# Headwaters fed by subterranean ice: potential climate refugia for mountain stream communities?

**DOI:** 10.1101/788273

**Authors:** Lusha M. Tronstad, Scott Hotaling, J. Joseph Giersch, Oliver J. Wilmot, Debra S. Finn

## Abstract

Near-term extirpations of macroinvertebrates are predicted for mountain streams worldwide as a warming climate drives the recession of high-elevation ice and snow. However, hydrological sources likely vary in their resistance to climate change and thus streams fed by more resistant sources could persist as climate refugia for imperiled biota. In 2015-2016, we measured habitat characteristics and quantified macroinvertebrate community structure along six alpine streams in the Teton Range, Wyoming, USA. Strong differences in habitat characteristics (e.g., temperature, bed stability, conductivity) confirmed three major stream sources: surface glaciers, perennial snowfields, and subterranean ice. Subterranean ice-fed streams – termed “icy seeps” – appear common in the Teton Range and elsewhere yet are globally understudied. Midges in the family Chironomidae dominated our study sites, representing 78.6% of all specimens sampled, with nematodes, caddisflies (*Neothremma*), and mayflies (*Epeorus*) also common. At the community-scale, glacier-and snowmelt-fed streams differed significantly in multivariate space, with icy-seep communities intermediate between them, incorporating components of both assemblages. Because the thermal environment of subterranean ice, including rock glaciers, is decoupled from large-scale climatic conditions, we predict that icy seeps will remain intact longer than streams fed by surface ice and snow. Furthermore, our results suggest that icy seeps are suitable habitat for many macroinvertebrates occupying streams fed by vulnerable hydrological sources. Thus, icy seeps may act as key climate refugia for mountain stream biodiversity, an idea in need of further investigation.

## INTRODUCTION

The highest rates of climate change are occurring above the permanent treeline in alpine and arctic ecosystems (Bradley et al. 2006). In the Rocky Mountains, warming is proceeding two to three times more quickly than the global average (Hansen et al. 2005, Pederson et al. 2010), resulting in extensive loss of glaciers and long-term snowpack (Hall & Fagre 2003). Streams fed by permanent ice may exhibit short-term increases in flow as air temperatures rise and source ice melting accelerates, but eventually they will shift to reduced flow and warm with the potential for intermittency or drying permanently (Hotaling et al. 2017). As climate change proceeds, invertebrate diversity at the mountain range scale is predicted to decrease due to both overall loss of habitat and summit traps, where the highest-altitude species and communities have nowhere left to disperse as warmer conditions and lower elevation communities shift upward. Biodiversity loss will be compounded by the loss of specific aquatic habitat types, particularly the unique conditions associated with meltwater from once-permanent hydrological sources like glaciers, snowfields, or subterranean ice (Brown et al. 2007, Milner et al. 2009, Jacobsen et al. 2012, Finn et al. 2013, 2014, Hotaling et al. 2017). Given that many alpine stream communities appear uniquely adapted to cold thermal regimes (but see Hotaling et al. 2019c), they are likely to be highly vulnerable to climate change as meltwater sources are lost (Giersch et al. 2017, Lencioni 2018). However, because alpine streams are heterogeneous with respect to hydrological source, potential also exists to identify stream types that may be locally buffered from broad-scale climate patterns and therefore could represent climate refugia for alpine stream biota (Morelli et al. 2016).

A major, long-term focus in alpine stream biology has been understanding the links between hydrological sources, the in-stream conditions they promote, and resident biotic communities (Ward 1994, Hotaling et al. 2017). According to primary hydrological source following Ward (1994), three types of alpine streams have historically been recognized: surface glacier-fed, snowmelt-fed, and groundwater-fed streams. A fourth, understudied stream type also exists – icy seeps – which are fed by subterranean ice (Hotaling et al. 2017, Hotaling et al. 2019a). In mountain ecosystems, the most common form of subterranean ice are rock glaciers, masses of debris-covered ice that act as conveyor belts moving fallen rock and other debris slowly downhill (Anderson et al. 2018; Jones et al. 2019). There may be more than 10,000 rock glaciers in the western United States (Johnson 2018) and they are similarly common worldwide (e.g., Lilleøren et al. 2011, Scotti et al. 2013, Charbonneau & Smith. 2018). In contrast, there are ∼1,250 surface glaciers and ∼3,750 perennial snowfields in the western United States (Fountain et al. 2017). Due to insulating debris cover, rock glaciers are largely decoupled from external conditions (e.g., warm summer air temperatures; Clark et al. 1994, Anderson et al. 2018, Knight et al. 2019), and should persist on the landscape longer than surface glaciers and snowfields. Icy seeps, the outflow of rock glaciers and similar ice features, may therefore act as climate refugia for cold-adapted stream biodiversity (Brighenti et al. 2019a,b; Hotaling et al. 2019a).

Like many mountain ranges worldwide, virtually nothing is known of alpine stream ecology and biodiversity in the Teton Range, a granite-dominated subrange of the Rocky Mountains. Previous studies of montane (2,000-3,000 m), but not alpine, streams revealed considerable macroinvertebrate diversity in the region’s higher elevation streams (Tronstad et al. 2016). Generally speaking, groundwater aquifers appear rare on the Teton Range massif, and thus groundwater-fed streams are also rare (L.M.T., personal observation). Groundwater-fed streams are considered the most resistant alpine stream type to warming because they are not directly influenced by surface ice (e.g., Milner et al. 2009, Jacobsen et al. 2012). With a paucity of groundwater-fed streams in the region, stream biodiversity in the Teton Range may be especially vulnerable to climate change.

In this study, we addressed two major objectives. First, we made the first assessment of alpine stream macroinvertebrate diversity in the Teton Range. Second, we explored associations between primary hydrological sources (surface glaciers, snowfields, and subterranean ice) and community structure. Specifically, we asked if benthic communities associated with icy seeps have substantial taxonomic overlap with communities linked to sources more vulnerable to climate change (e.g., glacier and snowfields). Our study provides an important first perspective on an urgent need in freshwater ecology (see Brighenti et al., 2019a); testing whether an underappreciated but globally common alpine stream type – icy seeps fed by subterranean ice – may act as key refugia for mountain aquatic biodiversity threatened by global change. Our results also provide new insight into the biodiversity of one of North America’s flagship protected areas, Grand Teton National Park, and neighboring wilderness areas.

## MATERIALS AND METHODS

### Study area

During the summers of 2015 and 2016 (26 July-10 August), we sampled six streams in the Teton Range of Grand Teton National Park and the adjacent Jedediah Smith Wilderness in northwestern Wyoming, USA (Figure 1; Table 1). Study streams were selected to span the breadth of alpine hydrological sources we have observed in the Teton Range and included two streams fed by surface glaciers (‘glacier-fed’ hereafter), two fed by subterranean ice (‘icy seep’ hereafter) and two fed by permanent snowpack (‘snowmelt-fed’). In 2015, we sampled both upstream (near the source) and downstream sites on each stream (Figure 1). On average, upper sites were 111 m higher in elevation and 690 m in stream distance from lower sites (Table 1). In 2016, we re-sampled the upper sites with the same methods to assess inter-annual variability. We focused on upper sites because they were as ‘true’ to primary hydrological source as possible while lower sites inherently reflected various degrees of mixing among sources. In both years, snow depth in the range was lower than average (152 cm in May, 1981-2010) with 2015 and 2016 at 63.3% and 80% of normal, respectively (Teton Pass, USDA SNOTEL).

**Table 1.** Key characteristics of our study streams and sites in the Teton Range, Wyoming, USA. Distances to source (*D*) are cumulative. Both elevation and *D* are in meters.

| Stream | Code | Site | Type | Lat., Long. | Elev. | $D$ |
| --- | --- | --- | --- | --- | --- | --- |
| Petersen Glacier | PG | Upper | Glacier-fed | 43.782, -110.846 | 2,922 | 51 |
|  |  | Lower | Glacier-fed | 43.785, -110.841 | 2,900 | 673 |
| Middle Teton | MT | Upper | Glacier-fed | 43.728, -110.795 | 2,955 | 178 |
|  |  | Lower | Glacier-fed | 43.725, -110.790 | 2,802 | 780 |
| South Cascade Creek | SCC | Upper | Icy seep | 43.722, -110.838 | 3,152 | 165 |
|  |  | Lower | Icy seep | 43.729, -110.837 | 2,943 | 1,051 |
| Wind Cave | WC | Upper | Icy seep | 43.666, -110.961 | 2,692 | 29 |
|  |  | Lower | Icy seep | 43.667, -110.955 | 2,564 | 144 |
| S. Fork Teton Creek | SFT | Upper | Snowmelt | 43.691, -110.843 | 2,987 | 1,227 |
|  |  | Lower | Snowmelt | 43.693, -110.859 | 2,881 | 2,757 |
| N. Fork Teton Creek | NFT | Upper | Snowmelt | 43.777, -110.860 | 2,955 | 9 |
|  |  | Lower | Snowmelt | 43.775, -110.861 | 2,910 | 393 |

**Figure 1.**
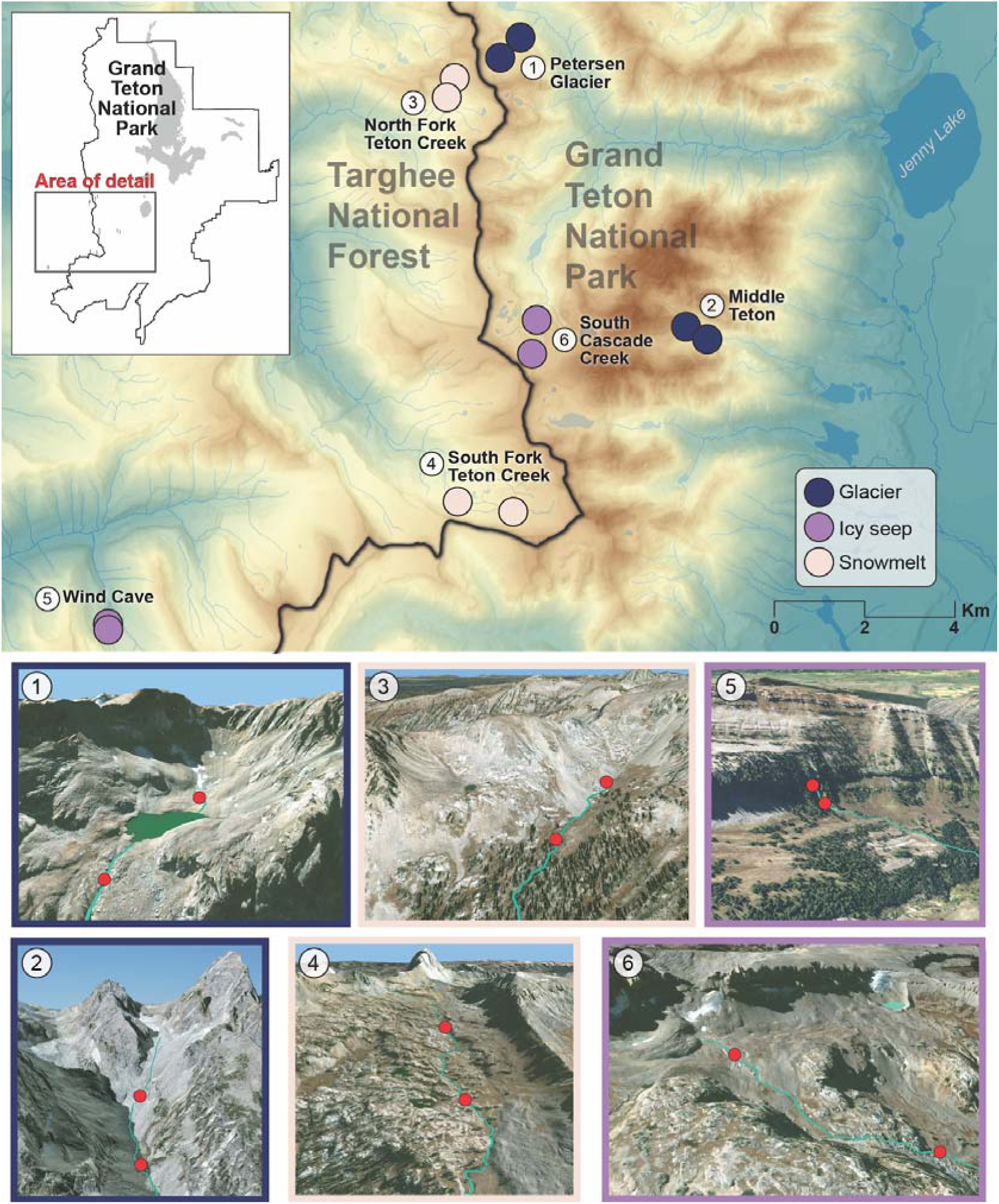
All streams and sites in the Teton Range, Wyoming, USA, included in this study. Upper sites were selected to be as close to the primary hydrological source as possible. Numbers in the top panel correspond to satellite imagery (DigitalGlobe, 15 October 2015, bottom panel) with stream channel (light blue) and sample sites (red dots) marked.

### Environmental data

At each site, we measured several environmental variables to characterize local habitat and evaluate whether in-stream environmental conditions varied among stream types. We measured water temperature for a full year (2015-2016) with *in situ* loggers (HOBO Pro v2, Onset Computer Corporation) that recorded temperature hourly. We measured specific conductivity (SPC), oxidation-reduction potential (ORP), pH, and dissolved oxygen (DO) with a Yellow Springs Instrument (YSI) Professional Plus multiparameter sonde calibrated at the trailhead (SPC, ORP, and pH) or at each site (DO). We estimated streambed stability with a modified version of the Pfankuch Index (PI) following Peckarsky et al. (2014). Total suspended sediments (TSS) were calculated by filtering known volumes of streamwater through pre-weighed filters (PALL Type A/E glass fiber filters) and measuring dry mass to the 10^−5^ grams. We used analysis of variance (ANOVA) statistical tests and the R package ‘plyr’ (R Core Development Team 2017; Wickham 2011) to characterize differences in environmental variables (Pfankuch Index, specific conductivity and total suspended solids) among stream types and between study sites. When stream type was significant (α = 0.05), we used Tukey’s HSD to distinguish which stream types differed from one another (*P* ≤ 0.05). Our use of Tukey’s HSD tests was highly conservative, given the small sample size of streams in this study.

Using the annual temperature data from each stream, we calculated mean temperature for the entire year (T_YEAR_), mean temperature between the summer solstice (21 June) and autumn equinox (22 September; T_SUMMER_), and maximum annual temperature range (T_RANGE_). We also estimated the date when seasonal snow covered (S_ON_) and uncovered (S_OFF_) each site by visually inspecting thermographs from our *in situ* temperature loggers to approximate the date when intraday thermal variation ceased and in-stream temperatures became constantly ∼0°C (S_ON_) or the opposite occurred (S_OFF_). An example thermograph indicating S_ON_ and S_OFF_ points is provided in Figure S1. Using S_ON_ and S_OFF_, we calculated S_DURATION_, the total days a site was snow covered between study years (S_DURATION_ = S_ON_ - S_OFF_). Finally, we used Principle Components Analysis (PCA; PC-ORD, McCune and Mefford 2006) to characterize the upper sites on each stream according to four variables (SPC, TSS, PI, T_RANGE_) which comprise a modified ‘glaciality index’ (Ilg & Castella 2006). Because T_SUMMER_ was correlated with T_RANGE_, we only included T_RANGE_ in the PCA. The glaciality index has been useful in characterizing alpine stream hydrological sources globally (e.g., Finn et al. 2013, Cauvy-Fraunié et al. 2015, Hotaling et al. 2019b).

### Macroinvertebrate sampling

We quantitatively sampled benthic macroinvertebrates at each site using a Surber sampler (Area: 0.09 m^2^, Mesh size: 243 μm). At each location, a composite sample of 5-10 replicates was collected depending on stream size, apparent biomass, and microhabitat diversity. Samples were elutriated in the field to reduce the amount of inorganic substrate and stored in Whirl-Pak bags with ∼80% ethanol. In the laboratory, invertebrate samples were divided into large (>2 mm mesh) and small (between 250 μm and 2 mm) fractions. For the large fraction, all invertebrates were identified. The small fraction was subsampled using the record player method when invertebrates were numerous (Waters 1969). Specimens were sorted, identified to the lowest taxonomic level possible using keys in Merritt et al. (2008) and Thorp & Covich (2010), and counted under a dissecting microscope. Insects were typically identified to genus when mature specimens were present except Chironomidae which were classified as either Tanypodinae or non-Tanypodinae. We estimated invertebrate density by summing the total number of individuals for a given site and dividing by the area of streambed sampled. We calculated biomass by measuring the length of the first 20 individuals of each taxon and then using length-mass regressions to estimate individual biomass (Benke et al. 1999). We multiplied the mean individual biomass for each taxon by the total number collected to estimate total biomass.

### Biological data analysis

We used ANOVAs and Tukey’s HSD tests performed on data summarized with ‘plyr’ (R Core Development Team 2017; Wickham 2011) to characterize differences in invertebrate density, biomass, and richness among stream types and study sites. To assess the relationship between taxonomic richness or biomass with stream characteristics of interest, namely snow cover (S_DURATION_), temperature (T_SUMMER_) and stability (Pfankuch Index), we performed both Pearson and Spearman’s rank-order correlations using the R package ‘Hmisc’ (Harrell Jr. 2013). For correlation analyses, we focused exclusively on upper sites and averaged taxonomic richness and biomass between 2015 and 2016.

We evaluated differences in community structure across streams, sites, and study years using non-metric multidimensional scaling (NMS) with PC-ORD (McCune & Mefford 2006). We log_10_ (*n* + 1) transformed density data for all taxa, removed rare taxa (either those private to a single site in the matrix and/or representing < 1% of the total abundance), and used Sørensen’s dissimilarities to create distance matrices. We ran NMS analyses independently on two data matrices: one including each of the upper and lower sites collected in 2015 only (N = 12 sites) and the other including only the upper sites (sampled in both 2015 and 2016; N = 12 sites). Dimensionality of the final solutions was chosen as the number of axes that produced the lowest stress following 200 iterations. Following NMS we applied multi-response permutation procedures (MRPP) in PC-ORD to assess whether there were differences in either community structure and/or mean community distance within the following groups: upstream versus downstream sites (2015 only) and among stream types for upper sites only (2015 and 2016). We then used Indicator Species Analysis (ISA) in PC-ORD to assess whether any specific taxa in the input matrices were indicative of (a) upper or lower sites or (b) a specific stream type. In ISA, the indicator value (IV) for a taxon is the test statistic. The maximum IV is 100, which indicates that a taxon is found in one group alone and is absent from other groups. We tested for significance of all IVs using Monte Carlo randomizations with 15,000 permutations of the data for each of the two tests (a and b above).

## RESULTS

### Environmental variation

Our upper sites clearly separated into three groups according to the glaciality index: glacier-fed streams, snowmelt-fed streams, and icy seeps (Figure 2; Table 2). Glacier-fed streams had less stable streambeds than both icy seeps and snowmelt-fed streams (F = 19.9, df = 2, *P* = 0.0001; Tukey’s HSD, *P* < 0.001; Table 2). Annual temperature range (T_RANGE_) was highest in snowmelt-fed streams and lowest in icy seeps. Specific conductivity was highest in icy seeps (SPC >100 μS cm^-1^ at upper sites; F = 30.6, df = 2, *P* < 0.001; Tukey’s HSD, *P* < 0.001) and lowest in glacier-fed streams (Table 2). More suspended solids (TSS) were present in glacier-fed streams (mean = 0.157 g/L) compared to snowmelt-fed streams (F = 3.6, df = 2, *P* = 0.44; Tukey’s HSD, *P* = 0.039; Table 2). Summer temperatures (T_SUMMER_) were lowest in glacier-fed streams (mean = 1.6 °C) and icy seeps (mean = 2.2 °C), and higher in snowmelt-fed streams (mean = 6.9 °C; Table 2). Upper sites were on average 1.2 °C colder in the summer and had less stable stream beds than lower sites (mean PI, upper = 27; mean PI, lower = 20.33; Table 1). Other environmental variables (i.e., DO, pH, ORP, days under snow) did not vary among stream types or between upper and lower sites (Table 2).

**Table 2.**
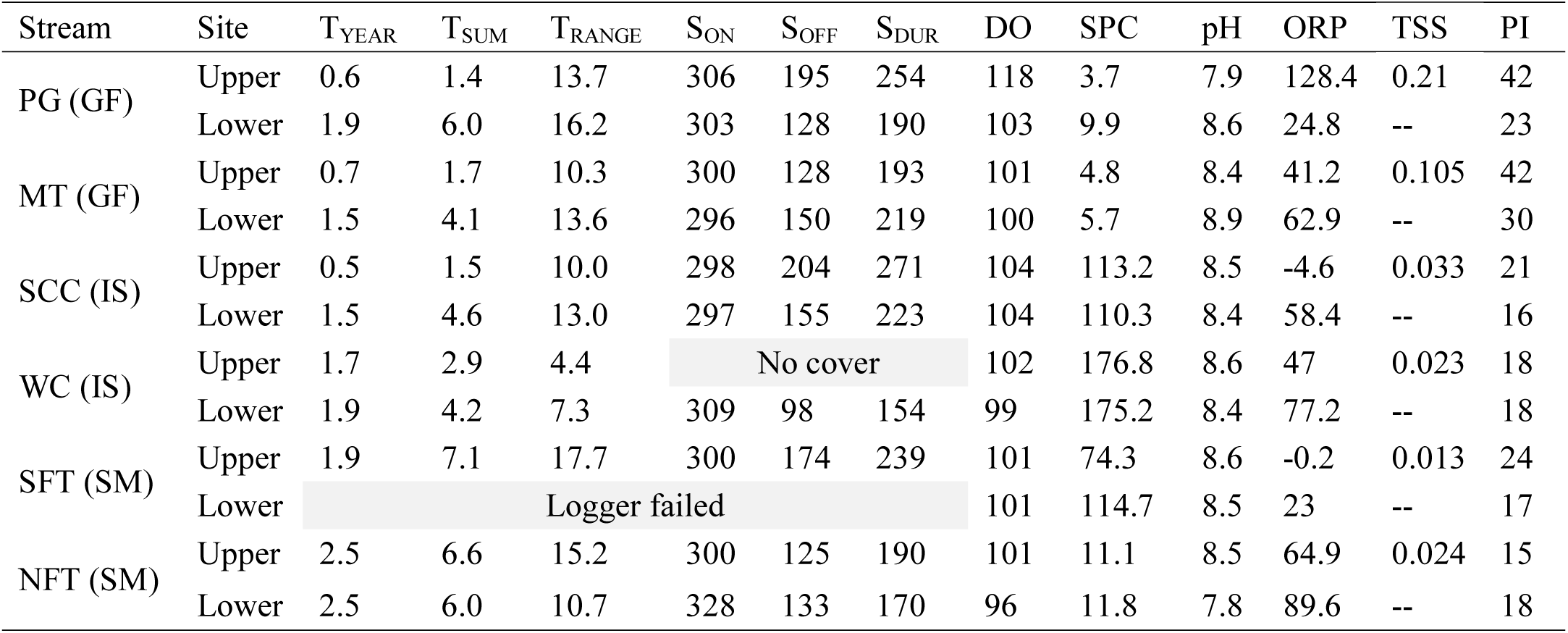
Environmental characteristics of the streams and sites included in this study. T_YEAR_: mean temperature for a calendar year. T_SUM_: mean temperature between the summer solstice (21 June) and autumn equinox (22 September). T_RANGE_: the difference between the maximum and minimum temperatures recorded in the stream during the calendar year. S_ON_ and S_OFF_ indicate the day of the year when a site was covered (S_ON_) or uncovered (S_OFF_) by snow according to thermographs. S_DURATION_ (S_DUR_ below) is the approximate total number of days each site spent under snow in 2015-2016. Abbreviations and units: temperature (T; °C), dissolved oxygen (DO, percent saturation), specific conductivity (SPC; μS cm^-1^), oxidation-reduction potential (ORP, mV), total suspended solids (TSS; g/L; upper sites only) and the Pfankuch Index (PI; higher values correspond to a less stable streambed). All data (except for S_ON_, S_OFF_, and S_DUR_) are for 2015 only. Stream codes are provided in Table 1 and parentheses after the stream name indicate the stream type (GF: glacier-fed; SM: snowmelt-fed; IS: icy seep).

| Stream | Site | $T_{\text{YEAR}}$ | $T_{\text{SUM}}$ | $T_{\text{RANGE}}$ | $S_{\text{ON}}$ | $S_{\text{OFF}}$ | $S_{\text{DUR}}$ | DO | SPC | pH | ORP | TSS | PI |
| --- | --- | --- | --- | --- | --- | --- | --- | --- | --- | --- | --- | --- | --- |
| PG (GF) | Upper | 0.6 | 1.4 | 13.7 | 306 | 195 | 254 | 118 | 3.7 | 7.9 | 128.4 | 0.21 | 42 |
|  | Lower | 1.9 | 6.0 | 16.2 | 303 | 128 | 190 | 103 | 9.9 | 8.6 | 24.8 | -- | 23 |
| MT (GF) | Upper | 0.7 | 1.7 | 10.3 | 300 | 128 | 193 | 101 | 4.8 | 8.4 | 41.2 | 0.105 | 42 |
|  | Lower | 1.5 | 4.1 | 13.6 | 296 | 150 | 219 | 100 | 5.7 | 8.9 | 62.9 | -- | 30 |
| SCC (IS) | Upper | 0.5 | 1.5 | 10.0 | 298 | 204 | 271 | 104 | 113.2 | 8.5 | -4.6 | 0.033 | 21 |
|  | Lower | 1.5 | 4.6 | 13.0 | 297 | 155 | 223 | 104 | 110.3 | 8.4 | 58.4 | -- | 16 |
| WC (IS) | Upper | 1.7 | 2.9 | 4.4 | No cover |  |  | 102 | 176.8 | 8.6 | 47 | 0.023 | 18 |
|  | Lower | 1.9 | 4.2 | 7.3 | 309 | 98 | 154 | 99 | 175.2 | 8.4 | 77.2 | -- | 18 |
| SFT (SM) | Upper | 1.9 | 7.1 | 17.7 | 300 | 174 | 239 | 101 | 74.3 | 8.6 | -0.2 | 0.013 | 24 |
|  | Lower | Logger failed |  |  |  |  |  | 101 | 114.7 | 8.5 | 23 | -- | 17 |
| NFT (SM) | Upper | 2.5 | 6.6 | 15.2 | 300 | 125 | 190 | 101 | 11.1 | 8.5 | 64.9 | 0.024 | 15 |
|  | Lower | 2.5 | 6.0 | 10.7 | 328 | 133 | 170 | 96 | 11.8 | 7.8 | 89.6 | -- | 18 |

**Figure 2.**
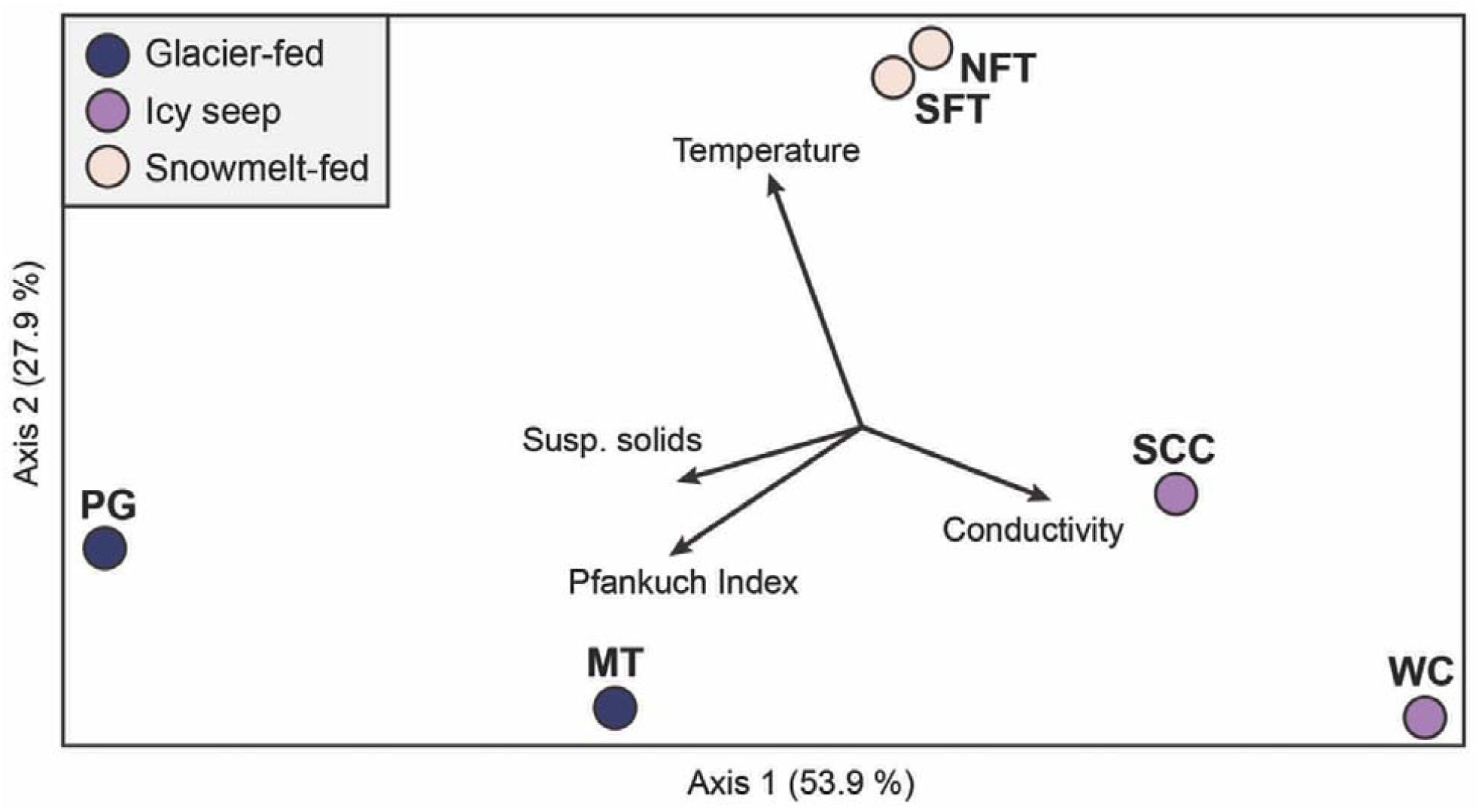
Principal components analysis (PCA) results showing environmental variation among the upper sites according to the four variables of the glaciality index [specific conductivity, streambed stability (Pfankuch Index), suspended solids, and temperature (T_RANGE_)].

### Biological variation

We collected 35 total invertebrate taxa of which 28 were insects (Supplementary Appendix 1). Insects composed 95% of the total mean densities and 92% of the total biomass. While invertebrate densities varied across stream types, biomass and richness were highest in snowmelt-fed streams (Figure 3A). At upper sites, total macroinvertebrate density did not differ among stream types (F = 1.3, df = 2, *P* = 0.31; Figure 3A), but biomass was ∼7x higher in snowmelt-fed streams compared to glacier-fed streams and icy seeps (F = 7.1, df = 2, *P* < 0.009; Tukey’s HSD, *P* ≤ 0.02; Figure 3B). Additionally, invertebrate richness was ∼2x higher in snowmelt-fed streams than glacier-fed streams and icy seeps (F = 10.4, df = 2, *P* < 0.002, Tukey’s HSD, *P* < 0.008; Figure 3C). We observed generally higher and more variable invertebrate densities (F = 2.7, df = 1, *P* = 0.13; Figure 3D) and higher biomass (F = 2.6, df = 1, *P* = 0.13; Figure 3E) at lower sites. We also observed ∼50% more taxa at lower sites (F = 8.5, df = 1, *P* = 0.012; Figure 3F). Only T_SUMMER_ was significantly correlated with richness or biomass (Richness: Pearson *r* = 0.81, *P* = 0.002; Biomass: Pearson *r* = 0.63, *P* = 0.038; Figure 4). Spearman’s rank-order correlations of the same relationships exhibited similar patterns, again with only T_SUMMER_ exhibiting significant correlations with richness (Spearman’s *r* = 0.89, *P* = 0.003) and biomass (Spearman’s *r* = 0.67, *P* = 0.023).

**Figure 3.**
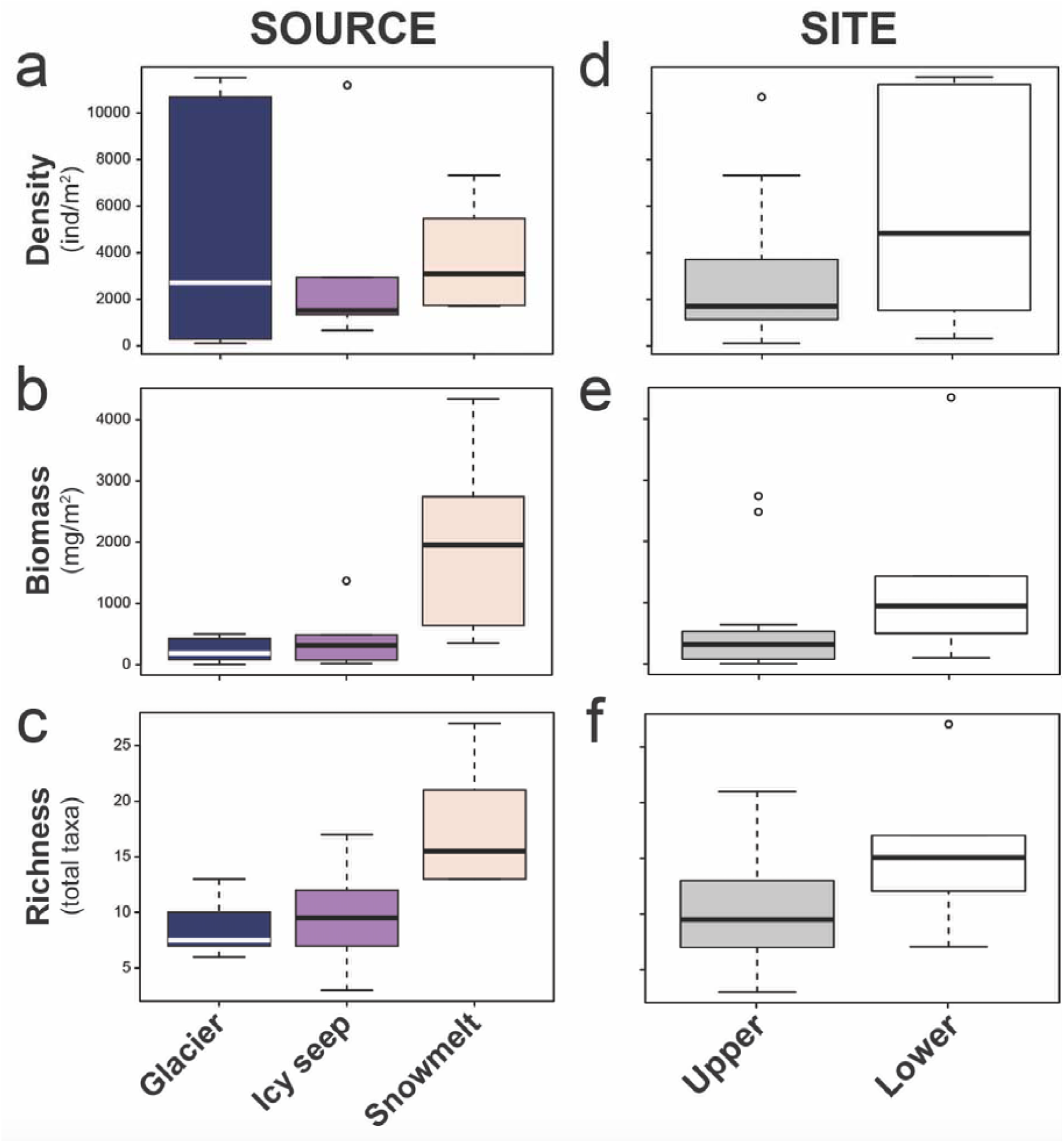
Macroinvertebrate density, biomass and richness among sites categorized by hydrologic source (a-c; upper sites only in 2015 and 2016) or site (d-f; 2015 and 2106 data for upper sites, 2015 data only for lower sites). Bold lines are median values, lower and upper box limits indicate the 25^th^ and 75^th^ percentiles, respectively, and whiskers represent the lower and upper limits of the data (excluding outliers which are shown as circles).

**Figure 4.**
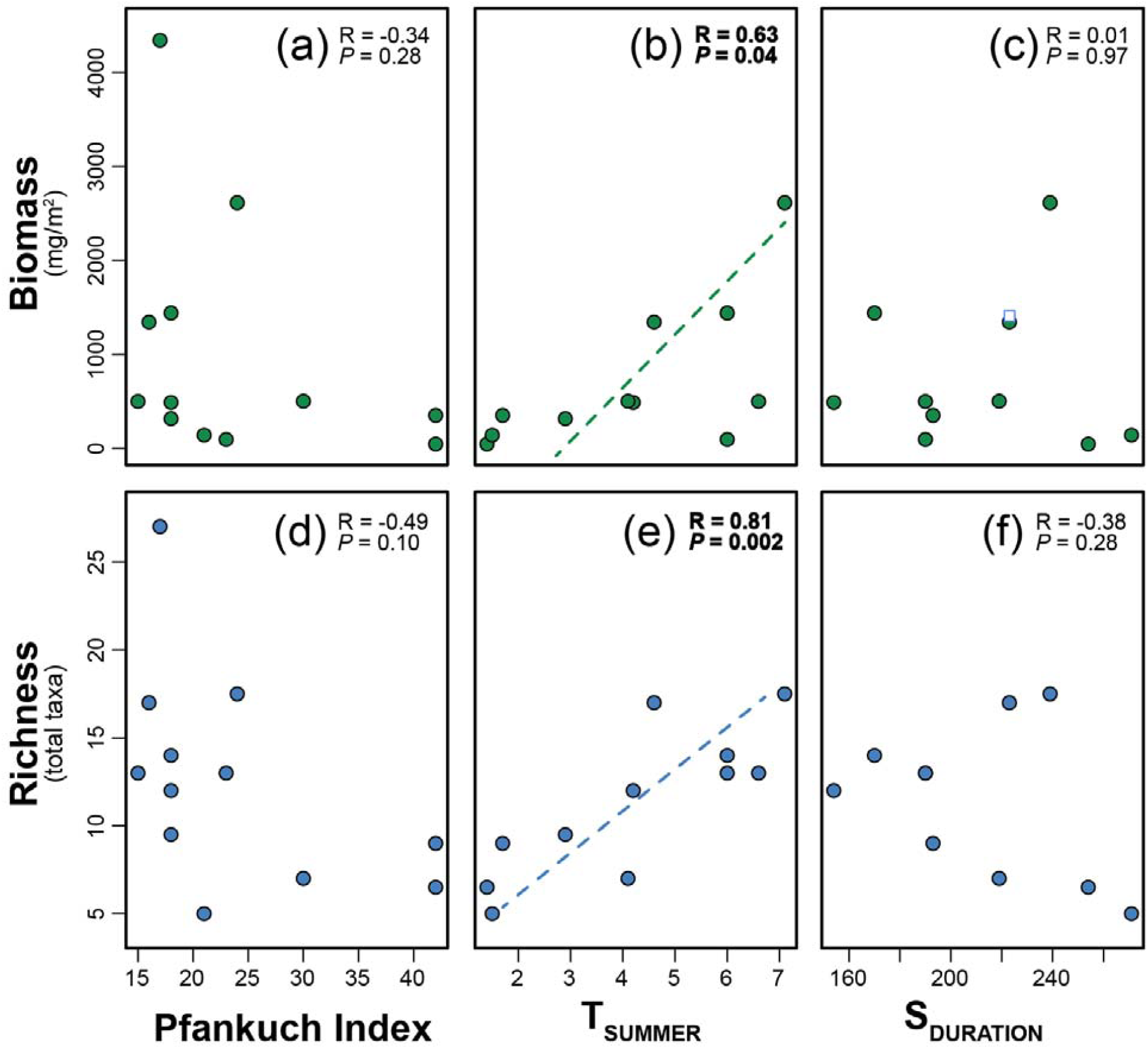
Bivariate plots of biomass (a-c) and richness (d-f) versus the Pfankuch Index, a measure of stream bed stability (higher = less stable), S_DURATION_ (the approximate number of days each stream was snow-covered in 2015-2016), and T_SUMMER_ (mean stream temperature between the summer and autumn solstices). Significant Pearson correlations (*P* < 0.05) are in bold with trendlines shown. The number of data points varies across plots due to a lack of snow cover at upper Wind Cave during the study period and a failed temperature logger at lower South Fork Teton Creek (see Table 2).

The most common invertebrates in high-elevation Teton streams were non-Tanypodinae midges (78.6% of all specimens sampled; >100 individuals/m^2^ at all sites; Appendix 1) followed by *Neothremma* (caddisflies), nematodes, and *Epeorus* (mayflies). Additionally, 45.7% of taxa had densities >10 individual/m^2^ among sites. Three taxa (Perlodidae, *Zapada* and Empididae) were present at 67% of sites and twelve taxa (34.3%), including *L. tetonica*, were only found at a single site (Appendix 1). Overall, we observed more stoneflies (6 taxa) than caddisflies (5 taxa) and mayflies (4 taxa; Appendix 1). Only two taxa (midges and Collembola) were collected at upper sites of both glacier-fed streams (Appendix 1). Two taxa were also present at both upper icy seep sites (midges and the stonefly genus, *Zapada*) and four taxa were present in upper snowmelt-fed sites [midges, stoneflies (Perlodidae), caddisflies (*Allomyia*), and flatworms (Turbellaria)]. Of those, only Turbellaria were present at consistently high numbers (e.g., >20 individuals/m^2^ per site; Appendix 1). No taxon exclusively occurred in one stream type and three occurred in upper sites of all stream types (midges, *Allomyia* caddisflies, and *Helodon* black flies; Appendix 1).

The most stable NMS solution comparing community structure between upper and lower sites was three-dimensional (stress = 9.6) with the first two axes explaining 88% of the total variation. Communities from the six lower sites overlapped substantially in ordination space with those from the six upper sites (MRPP *A* = 0.013; *P* = 0.27). However, the mean pairwise community distance was greater among upper versus lower sites (0.58 vs. 0.47), a trend that is apparent in the NMS bi-plot (Figure 5A). Indicator species analyses revealed no taxon with IV scores approaching the maximum of 100 (Tables S2-S3), an expected result when two assemblages substantially overlap in ordination space. Mean IV across all taxa was 44.3, and Empididae had the maximum IV (78.9; *P* = 0.025) to lower sites.

**Figure 5.**
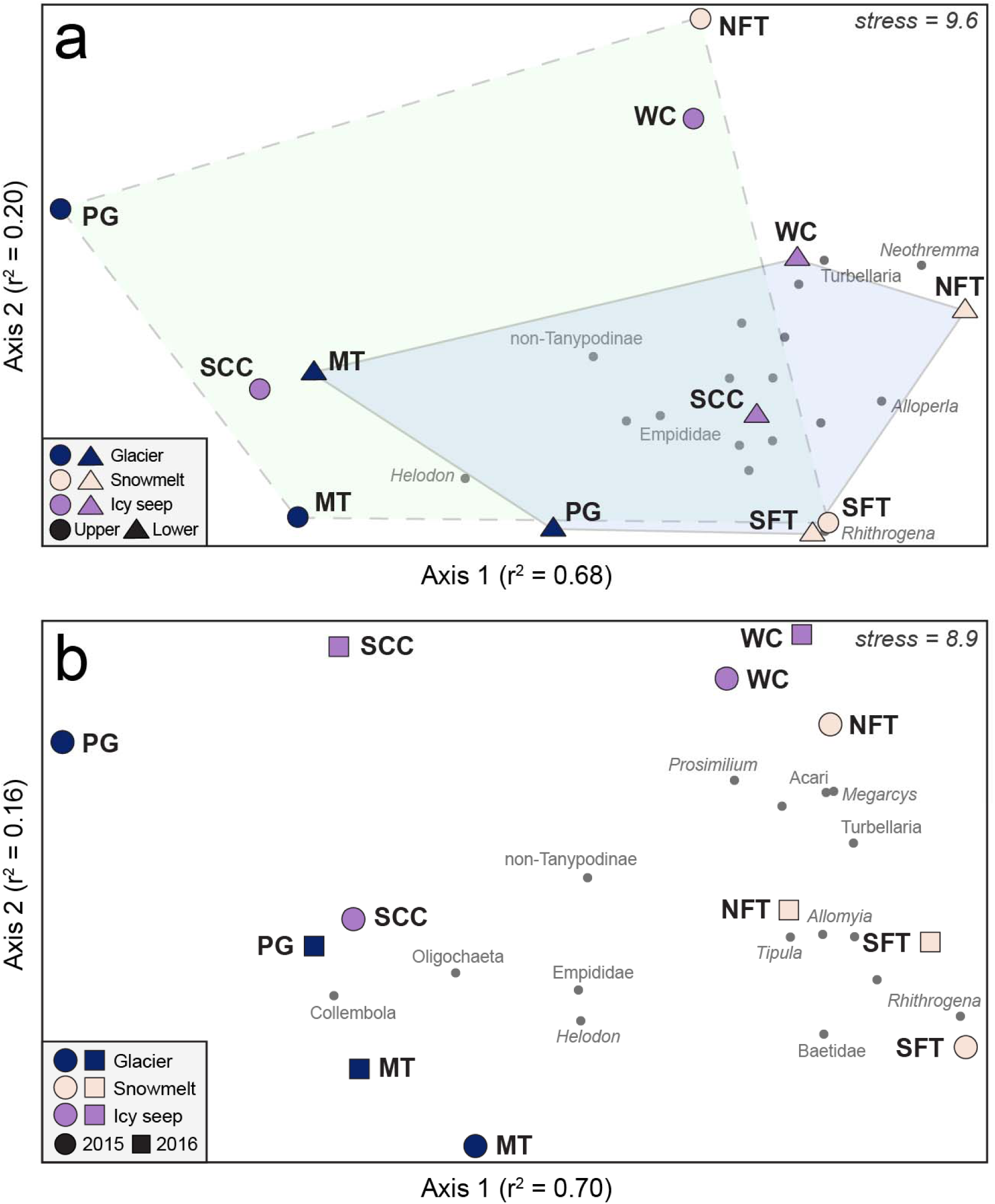
Non-metric multidimensional scaling (NMS) plots of macroinvertebrate communities comparing (a) upper (circles) versus lower (triangles) sites in 2015 and (b) upper sites only between 2015 (circles) and 2016 (squares). Colored polygons in (a) reflect the breadth of NMS space occupied by upper (dashed lines, green fill) and lower (solid lines, blue fill) sites.

The NMS analysis that included upper sites only (2015 and 2016) converged on a two-dimensional solution as the most stable result (stress = 8.9). The two axes explained 86% of the variation, with axis-1 explaining 70% alone. In general, communities occupying glacier-fed streams had the lowest axis-1 values, communities in snowmelt-fed streams had the highest axis-1 values and icy seep communities had intermediate axis-1 values (Figure 5B). MRPP results suggested that communities occupying these three stream types were significantly different from one another (*A* = 0.19; *P* = 0.006). Pairwise differences were strong between glacier-fed and snowmelt-fed communities (*A* = 0.21; *P* = 0.005) and were weaker but significant for the two pairs that included icy seep communities (icy seep vs. glacier-fed: *A* = 0.11; *P* = 0.05; icy seep vs. snowmelt: *A* = 0.12; *P* = 0.03). Indicator species analysis (Table S3) revealed IV scores >80 for three taxa, of which one was indicative of glacier-fed streams (Collembola; IV = 85.0; *P* = 0.02) and two were indicative of snowmelt streams (Tipulidae, IV = 82.1, *P* = 0.004; *Allomyia*, IV = 81.5, *P* = 0.004). Turbellaria were also weakly indicative of snowmelt streams (IV = 72.5; *P* = 0.011).

## DISCUSSION

As climate change proceeds and mountain glaciers recede, there is a need to develop a clearer understanding of how patterns of extant biodiversity and habitat heterogeneity are linked in high-elevation ecosystems. In this study, we provide the first description of macroinvertebrate diversity in the high Teton Range, Wyoming, where three major alpine stream types exist: glacier-fed streams, snowmelt-fed streams, and icy seeps. To the best of our knowledge, no streams fed by groundwater aquifers have been documented in the alpine zone of the granitic Teton Range. From a global perspective, icy seeps, which are fed by subterranean ice (primarily rock glaciers), rather than aquifers of liquid water, are of particular interest as they are likely to persist on the landscape longer than surface ice features (Hotaling et al. 2019a). For the Teton Range, a scarcity of high-elevation, groundwater-fed streams suggests that biodiversity in the region may be even more reliant on meltwater than similar mountain regions (e.g., European ranges, Brown et al. 2007; tropical Andes, Finn et al. 2016; Glacier National Park, Giersch et al. 2017). Thus, icy seep-associated climate refugia may be particularly important to Teton alpine stream biodiversity as climate change proceeds. In the streams sampled for this study, glacier-and snowmelt-fed streams exhibited significantly different invertebrate communities; however, icy seeps were intermediate between the two in terms of community structure and invertebrate density, biomass, and richness. These results suggest that icy seep communities share some characteristics with both glacier-and snowmelt-fed streams and have the potential to act as climate refugia for at least a subset of the unique communities present in each of the more vulnerable stream types. Thus, the potential for icy seeps and ice-influenced terrestrial refugia (Millar et al. 2015) to buffer climate-induced biodiversity loss has profound, global implications.

The recession of meltwater sources is predicted to strongly affect downstream invertebrate communities (Jacobsen et al. 2012). In the near term, rising in-stream temperatures are expected as ice melt comprises ever smaller proportions of stream flow. In alpine streams worldwide, warmer conditions have been correlated with increased species richness for microbial diversity (e.g., Wilhelm et al. 2013, Hotaling et al. 2019a), diatoms (Fell et al. 2018), and macroinvertebrates (Finn & Poff 2005, Jacobsen et al. 2012). Our study adds another line of evidence to this global pattern as we detected a positive correlation between species richness and mean summer temperature (T_SUMMER_). We also observed greater richness at lower (mean = 15 taxa) versus upper (mean = 10 taxa) sites which aligns with, and extends, the conclusions of Tronstad et al. (2016), the only other study to investigate longitudinal patterns of macroinvertebrate richness in montane streams of the Teton Range. Indeed, we observed far fewer taxa at our highest elevation sites (10 taxa at ∼3,150 m) versus the highest elevation sites included in Tronstad et al. (2016): 26 taxa at ∼2,700 m.

Although local (alpha) diversity will likely increase with warming water temperatures, among-stream (beta) diversity may decrease as more diverse, generalist communities shift upstream and specialized cold-adapted taxa are lost, effectively homogenizing biological diversity at the regional scale (Jacobsen et al. 2012, Wilhelm et al. 2013, Hotaling et al. 2017). In the Teton Range, we observed greater beta diversity at upper versus lower sites, and ISA revealed indicator taxa for snowmelt and glacier-fed streams, but not for icy seeps. Collectively, these patterns suggest that in the Teton Range, like elsewhere in the world, heterogeneous hydrological sources bolster regional-scale alpine stream biodiversity. Icy seep communities also appeared to bridge the taxonomic gap between glacier-and snowmelt-fed streams, and this pattern remained stable between years (Figure 5B). While clear patterns of community dissimilarity existed among our sites, ascribing a specific driver to these differences is difficult. Many factors likely shape community structure in headwater streams (and our estimation of it), including the primary source and its associated environmental regime, geographic location, and both the method and timing of sampling (e.g., a site recently uncovered from seasonal snowpack is likely to have a distinct community when compared to a site that melted out much earlier).

Our study represents the first perspective of macroinvertebrate biodiversity in the highest elevation streams of the Teton Range. As such, there are many areas where future study will expand, and refine, our conclusions. First, we did not have the resources to identify midges (family Chironomidae) to lower taxonomic resolution. Midges are diverse in alpine streams around the world (e.g., Montagna et al. 2016), and are typically the most common taxon (a pattern we also observed). Thus, the limited taxonomic resolution of midges in our study may have influenced our conclusions. For instance, Finn & Poff (2011) found 22 midge species in snowmelt-fed alpine streams of the Colorado Rockies. Given our focus on three stream types, we might expect even greater midge species diversity in the Teton Range. Second, our focus on a limited number of sites, lack of within-year temporal sampling, and use of a single sampling method/microhabitat (Surber samples from riffle habitats) rather than many approaches/habitats (e.g., Ghani et al. 2016, Tronstad & Hotaling 2017), limits our ability to confidently describe the full invertebrate community at a given site. We hope to fill these gaps in the future.

Ultimately, the degree to which alpine streams will be affected by climate change in terms of flow magnitude and persistence remains largely unknown. In general, studies of alpine stream ecology operate under the assumption that perennial flow will continue in the decades to come (e.g., Jacobsen et al. 2012), but this may not be the case (e.g., Haldorsen & Heim 1999, Herbst et al. 2019, Siebers et al. 2019). Thus, the biological ramifications of declining meltwater sources in places like the Teton Range, where groundwater aquifer-fed streams are scarce, may be even more profound than in ranges with a more equitable distribution of alpine stream sources (e.g., Glacier National Park, Giersch et al. 2017). Indeed, if alpine streams supported by surface glaciers and permanent snowfields transition to intermittency or dry completely, the future of biodiversity in these ecosystems may depend almost exclusively on icy seeps. Our study paired with the broader glaciological literature suggests there is room for optimism. Icy seeps have the potential to span a wide beta diversity profile, perhaps bridging the taxonomic gap between glacier-and snowmelt-fed communities (e.g., as observed for microbial diversity, Hotaling et al. 2019a). Moreover, rock glaciers and other subterranean ice forms (e.g., periglacial taluses, Millar et al. 2013) are common in alpine regions worldwide and likely the most resistant ice form to future warming. Clearly, research focused on subterranean ice sources and associated icy seeps, and specifically the biological communities and hydrological flows they support, represent a pressing need in alpine stream biology. We suggest that future studies incorporate temporal monitoring of multiple alpine stream types, including icy seeps, to separate year-to-year variation from true temporal signals. Long-term data will also allow for explicit tests of how biodiversity and habitat characteristics may be altered among alpine stream types as climate change proceeds.

## ACKNOWLEDGEMENTS

Financial support was provided by The University of Wyoming-National Park Service Research Station and the Teton Conservation District. We thank Lydia Zeglin for assistance in the field and Hanna Foster, Logan Fox, Alexis Lester, Jackson Marr, Jake Ruthven, Joe Wannemuehler, and Kara Wise for their help sorting invertebrate samples. Matthew Green provided helpful comments on the manuscript. Any use of trade, firm, or product names is for descriptive purposes only and does not imply endorsement by the U.S. Government.

