## Supplementary Materials for "Headwaters fed by subterranean ice: potential climate refugia for mountain stream communities?"

**TABLES**

**Supplementary Table 1.** Macroinvertebrate density (individuals/m^2^), biomass (mg/m^2^), and richness for each combination of site and year included in this study.

| Stream | Site | Year | Density | Biomass | Richness |
| --- | --- | --- | --- | --- | --- |
| Petersen Glacier | Lower | 2015 | 294 | 94 | 13 |
|  | Upper | 2015 | 116 | 5.2 | 6 |
|  | Upper | 2016 | 934 | 83 | 7 |
| Middle Teton | Lower | 2015 | 11,523 | 501 | 7 |
|  | Upper | 2015 | 10,694 | 274 | 8 |
|  | Upper | 2016 | 4,491 | 425 | 10 |
| South Cascade Creek | Lower | 2015 | 11,198 | 1,342 | 17 |
|  | Upper | 2015 | 1,545 | 77 | 7 |
|  | Upper | 2016 | 667 | 201 | 3 |
| Wind Cave | Lower | 2015 | 1,497 | 488 | 12 |
|  | Upper | 2015 | 1,346 | 211 | 9 |
|  | Upper | 2016 | 2,945 | 416 | 10 |
| South Fork Teton Creek | Lower | 2015 | 4,165 | 4,341 | 27 |
|  | Upper | 2015 | 2,041 | 2,483 | 14 |
|  | Upper | 2016 | 1,738 | 2,742 | 21 |
| North Fork Teton Creek | Lower | 2015 | 4,381 | 1,439 | 14 |
|  | Upper | 2015 | 1,696 | 355 | 13 |
|  | Upper | 2016 | 7,320 | 641 | 13 |

**Supplementary Table 2.** Results of an Indicator Species Analysis assessing the degree to which an individual taxon is associated with a given site (upper or lower; 2015 data only). A higher observed Indicator Value (IV) indicates stronger assignment to a group. Randomized IV and SD are the IVs and standard deviation, respectively, for randomized groups.

| Taxon | Group | Observed IV | Randomized IV | SD | *P* |
| --- | --- | --- | --- | --- | --- |
| Empididae | Lower | 78.9 | 47.3 | 11.6 | 0.025 |
| Perlodidae | Lower | 66.6 | 50.1 | 9.2 | 0.103 |
| Baetidae | Lower | 63 | 43.7 | 10.7 | 0.037 |
| *Prosimulium* | Lower | 62 | 40 | 12.8 | 0.067 |
| *Zapada* | Lower | 57 | 46.6 | 11.2 | 0.142 |
| *Rhyacophila* | Lower | 52.7 | 45.2 | 13.3 | 0.277 |
| non-Tanypodinae | Lower | 52.1 | 52.5 | 1.9 | 0.510 |
| Acari | Lower | 48.7 | 39.5 | 12.8 | 0.177 |
| *Epeorus* | Lower | 46.4 | 39.7 | 12.7 | 0.273 |
| Collembola | Lower | 45.7 | 43.6 | 11.5 | 0.324 |
| *Alloperla* | Lower | 41.1 | 31.4 | 12.8 | 0.305 |
| Nematoda | Lower | 33.3 | 22.4 | 10.0 | 0.456 |
| *Allomyia* | Upper | 29.8 | 43.7 | 11.5 | 1.000 |
| *Helodon* | Upper | 28.9 | 39.8 | 12.4 | 0.782 |
| Turbellaria | Upper | 24.4 | 36 | 11.8 | 1.000 |
| *Neothremma* | Lower | 13.5 | 29.4 | 12.1 | 1.000 |
| *Rhithrogena* | Lower | 8.5 | 19.8 | 12.4 | 1.000 |

**Supplementary Table 3.** Results of an Indicator Species Analysis assessing the degree to which an individual taxon is associated with a given stream type (glacier-fed, snowmelt-fed, or an icy seep; 2015 and 2016 data). A higher observed Indicator Value (IV) indicates stronger assignment to a group. Randomized IV and SD are the IVs and standard deviation, respectively, for randomized groups.

| Taxon | Group | Observed IV | Randomized IV | SD | *P* |
| --- | --- | --- | --- | --- | --- |
| Collembola | Glacier-fed | 85 | 35.3 | 14.6 | 0.019 |
| *Tipula* | Snowmelt-fed | 82.1 | 36.4 | 13.0 | 0.004 |
| *Allomyia* | Snowmelt-fed | 81.5 | 36.4 | 13.2 | 0.004 |
| Turbellaria | Snowmelt-fed | 72.5 | 36.1 | 12.8 | 0.011 |
| *Megarcys* | Snowmelt-fed | 61.2 | 36 | 12.9 | 0.078 |
| *Zapada* | Icy seep | 59.7 | 36.7 | 13.4 | 0.071 |
| *Rhyacophila* | Snowmelt-fed | 56.6 | 38.3 | 14.3 | 0.103 |
| Acari | Snowmelt-fed | 50.8 | 33.1 | 14.3 | 0.068 |
| *Rhithrogena* | Snowmelt-fed | 50 | 27 | 14.4 | 0.280 |
| *Epeorus* | Snowmelt-fed | 45.6 | 30.1 | 16.0 | 0.280 |
| *Helodon* | Glacier-fed | 38.8 | 38.6 | 11.4 | 0.363 |
| Baetidae | Snowmelt-fed | 36.7 | 28.9 | 15.6 | 0.280 |
| non-Tanypodinae | Glacier-fed | 33.8 | 35.9 | 1.3 | 0.976 |
| Oligochaeta | Glacier-fed | 26.3 | 32.8 | 14.7 | 0.710 |
| *Prosimulium* | Snowmelt-fed | 16.5 | 30.9 | 16.1 | 1.000 |
| Empididae | Snowmelt-fed | 10.6 | 29 | 15.2 | 1.000 |

**FIGURES**

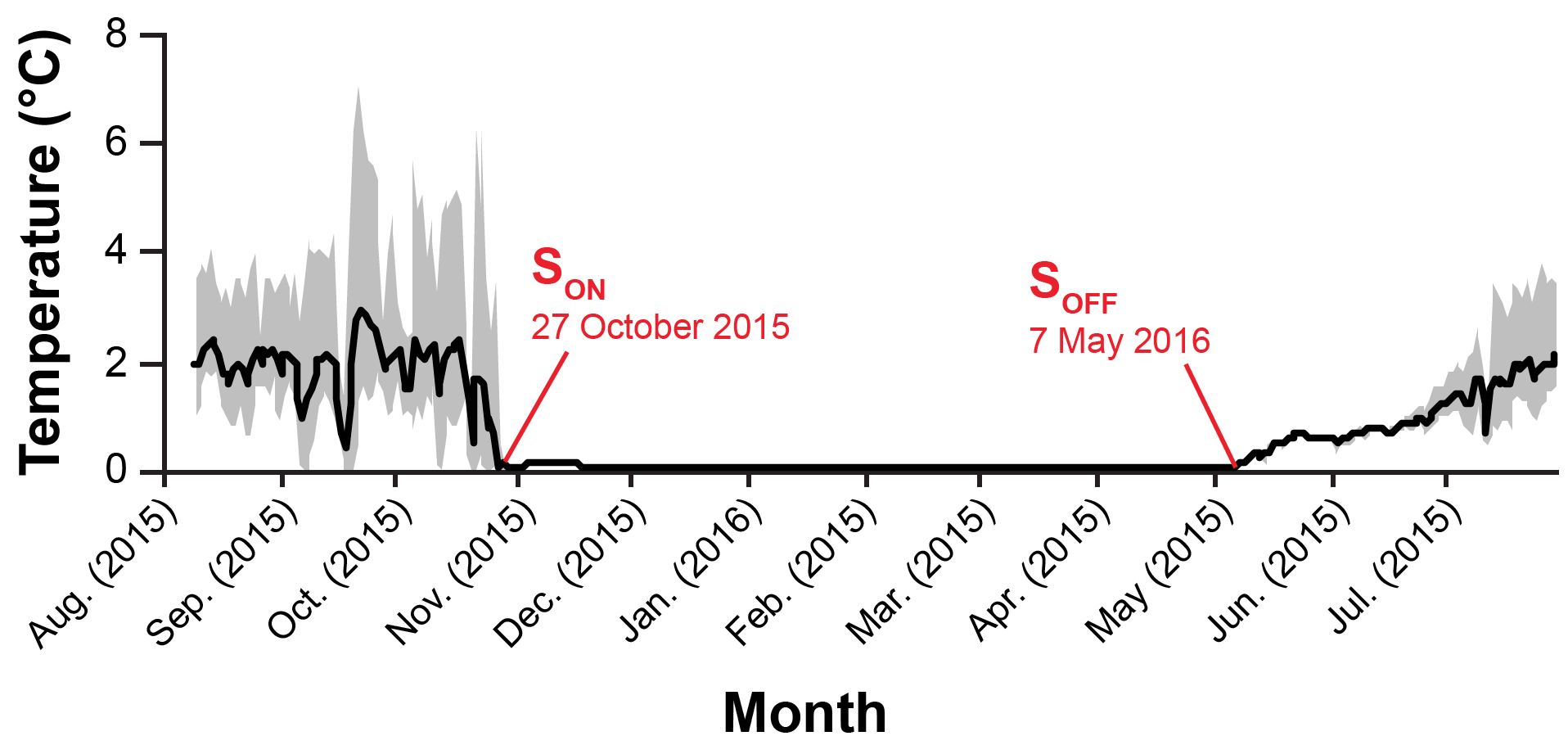

**Figure S1.** An example thermograph of a stream (Middle Teton, Upper) included in this study showing the dates when the stream became covered (SON) and uncovered (SOFF) by snow. The dark line is the mean temperature for each day and the gray area is the intraday thermal variation (minimum to maximum temperatures observed).

**APPENDICES**

**Supplementary Appendix 1.** All taxa observed across study sites in 2015 when both upper and lower sites were sampled. The first two letters in each column represent site names with U or L indicating upper or lower sites, respectively. Densities (ind/m^2^) for each taxon are given. MT: Middle Teton, NFT: North Fork Teton Creek, PG: Petersen Glacier, SCC: South Cascade Canyon, SFT: South Fork Teton Creek, WC: Wind Cave.

| Taxon | MTU­ | MTL | NFTU | NFTL | PGU | PGL | SCCU | SCCL | SFTU | SFTL | WCU | WCL |
| --- | --- | --- | --- | --- | --- | --- | --- | --- | --- | --- | --- | --- |
| **Non-insect invertebrates** | | | | | | | | | | | | |
| Acari | 0 | 0 | 14 | 29 | 0 | 3 | 0 | 29 | 0 | 56 | 5 | 0 |
| Collembola | 4 | 0 | 0 | 14 | 2 | 2 | 6 | 29 | 0 | 18 | 0 | 0 |
| Nematoda | 0 | 0 | 0 | 0 | 0 | 0 | 0 | 1521 | 0 | 6 | 0 | 0 |
| Oligochaeta | 0 | 0 | 0 | 14 | 2 | 0 | 22 | 0 | 0 | 6 | 0 | 14 |
| Ostracoda | 0 | 0 | 0 | 72 | 0 | 0 | 0 | 0 | 0 | 0 | 0 | 0 |
| Sphaeriidae | 0 | 0 | 0 | 0 | 0 | 0 | 0 | 0 | 0 | 2 | 0 | 0 |
| Turbellaria | 0 | 0 | 29 | 660 | 0 | 0 | 0 | 0 | 43 | 0 | 40 | 140 |
| **Insects** |  |  |  |  |  |  |  |  |  |  |  |  |
| **Coleoptera** |  |  |  |  |  |  |  |  |  |  |  |  |
| Elmidae | 0 | 0 | 14 | 0 | 0 | 0 | 0 | 0 | 0 | 0 | 0 | 0 |
| **Diptera** |  |  |  |  |  |  |  |  |  |  |  |  |
| *Clinocera* | 7 | 4 | 0 | 0 | 0 | 2 | 4 | 22 | 0 | 4 | 0 | 1 |
| *Dicranota* | 0 | 4 | 7 | 0 | 0 | 0 | 0 | 0 | 0 | 15 | 0 | 0 |
| Empididae | 7 | 4 | 0 | 43 | 0 | 2 | 4 | 79 | 0 | 8 | 0 | 1 |
| *Helodon* | 424 | 60 | 0 | 0 | 0 | 8 | 13 | 0 | 22 | 9 | 0 | 0 |
| non-Tanypodinae | 10237 | 11445 | 1298 | 1464 | 108 | 117 | 1496 | 9078 | 481 | 2919 | 1192 | 945 |
| *Oreogeton* | 0 | 0 | 0 | 43 | 0 | 0 | 0 | 0 | 0 | 0 | 0 | 0 |
| *Probezzia* | 0 | 0 | 0 | 0 | 0 | 0 | 0 | 0 | 0 | 28 | 0 | 0 |
| *Prosimulium* | 0 | 0 | 186 | 244 | 0 | 3 | 0 | 65 | 0 | 4 | 0 | 11 |
| *Simulium* | 0 | 0 | 0 | 0 | 0 | 3 | 0 | 0 | 0 | 0 | 0 | 0 |
| *Tipula* | 0 | 0 | 0 | 0 | 0 | 0 | 0 | 2 | 29 | 2 | 0 | 0 |
| **Ephemeroptera** |  |  |  |  |  |  |  |  |  |  |  |  |
| Baetidae | 14 | 0 | 0 | 72 | 0 | 50 | 0 | 58 | 36 | 82 | 0 | 16 |
| *Drunella* | 0 | 0 | 0 | 0 | 0 | 0 | 0 | 0 | 0 | 95 | 0 | 0 |
| *Epeorus* | 0 | 0 | 0 | 14 | 2 | 81 | 0 | 201 | 890 | 292 | 0 | 0 |
| *Rhithrogena* | 0 | 0 | 0 | 0 | 0 | 0 | 0 | 0 | 100 | 118 | 0 | 0 |
| **Hemiptera** |  |  |  |  |  |  |  |  |  |  |  |  |
| *Lipogomphus* | 0 | 0 | 0 | 0 | 0 | 0 | 0 | 0 | 0 | 4 | 0 | 0 |
| *Pentacora* | 0 | 0 | 0 | 0 | 2 | 0 | 0 | 0 | 0 | 0 | 0 | 0 |
| **Plecoptera** |  |  |  |  |  |  |  |  |  |  |  |  |
| *Alloperla* | 0 | 0 | 0 | 115 | 0 | 0 | 0 | 0 | 7 | 20 | 0 | 5 |
| *Lednia tetonica* | 4 | 0 | 0 | 0 | 0 | 0 | 0 | 0 | 0 | 0 | 0 | 0 |
| Leuctridae | 0 | 0 | 0 | 71 | 0 | 0 | 0 | 0 | 0 | 0 | 0 | 0 |
| *Megarcys* | 0 | 0 | 108 | 29 | 0 | 0 | 0 | 0 | 9 | 77 | 48 | 89 |
| Perlodidae | 0 | 9 | 108 | 29 | 0 | 8 | 0 | 43 | 9 | 249 | 48 | 89 |
| *Zapada* | 0 | 0 | 0 | 210 | 0 | 11 | 4 | 55 | 118 | 194 | 58 | 263 |
| **Trichoptera** |  |  |  |  |  |  |  |  |  |  |  |  |
| *Allomyia* | 4 | 0 | 5 | 0 | 0 | 0 | 0 | 18 | 11 | 13 | 1 | 12 |
| Limnephilidae | 0 | 0 | 5 | 0 | 0 | 0 | 0 | 0 | 0 | 0 | 0 | 0 |
| *Neothremma* | 0 | 0 | 2 | 2368 | 0 | 0 | 0 | 0 | 0 | 0 | 1 | 0 |
| *Parapsyche* | 0 | 0 | 0 | 0 | 0 | 0 | 0 | 0 | 2 | 2 | 0 | 0 |
| *Rhyacophila* | 0 | 2 | 0 | 80 | 0 | 0 | 0 | 2 | 185 | 17 | 1 | 1 |
